# Clonal memory in human embryonic stem cells biases fate potential during endoderm differentiation

**DOI:** 10.64898/2026.09.02.748863

**Authors:** Madeleine Linneberg-Agerholm, Álvaro Bustos Gutiérrez, Stine Lind Hansen, Yutaka Shimizu, Nicolaj Strøyer Christophersen, Samantha A. Morris, Atefeh Lafzi

**Author notes:** Authors contributed equally (Álvaro Bustos Gutiérrez, Stine Lind Hansen).

## Abstract

Cell fate decisions during development are shaped not only by extrinsic signals but also by heritable intrinsic states passed on across cell division. The extent to which this phenomenon, termed clonal memory, can explain the persistent heterogeneity observed from directed differentiation of human embryonic stem cells is unclear. Here, we combine lineage tracing with single-cell transcriptomics and chromatin accessibility profiling to track clonal behaviour across human embryonic stem cell differentiation towards definitive endoderm. Using a lentiviral barcoding system coupled with a split-well sampling strategy, we find that clonally related cells exhibit reproducible, probabilistic fate outcomes that cannot be explained by signalling environment alone. Fate-biased clones are transcriptionally indistinguishable at the pluripotent stage yet display distinct chromatin accessibility landscapes at lineage-specific cis-regulatory elements. Pre-existing accessibility at these lineage-specific regulatory regions distinguish clones that undergo successful endoderm differentiation from those that generate off-target mesoderm derivatives. Together, these findings provide an explanation for how off-target populations arise during directed differentiation, identifying heritable chromatin states within pluripotent cultures as a source of variability relevant to stem cell-derived *in vitro* models and cell therapies.

## Introduction

Human embryonic stem cells (hESCs) possess the remarkable ability to undergo self-renewal and differentiate into all cell types of the body, making them a valuable resource for understanding human development, disease modelling and regenerative medicine. The capacity of hESCs for directed differentiation to clinically relevant cell types has positioned pluripotent stem cell-based therapies as an area of vast potential for treating chronic and progressive diseases^1^. Despite decades of optimising both pluripotent stem cell culture conditions and directed differentiation protocols, a persistent challenge remains that pluripotent stem cells, based on expression of canonical markers such as NANOG, OCT4 and SOX2^2^, continue to produce heterogenous outcomes in response to identical signalling cues^3^. Understanding the sources and consequences of this heterogeneity is fundamental to realizing the full potential of stem cell-based therapies and cellular models.

Single-cell transcriptomics has led to new opportunities for characterising heterogeneity in hESC populations^4–6^, capturing individual cells displaying variable gene expression dynamics even under uniform culture conditions. However, whether this transcriptional variability represents transient stochastic fluctuations or stable, heritable cell states that may bias differentiation outcomes remains less clear. Clonal lineage tracing paired with single-cell RNA-sequencing (scRNA-seq) offers a powerful approach to dissect the origin of cellular heterogeneity by simultaneously capturing the progeny of individual cells together with the entire transcriptome^7–9^. Emerging evidence from various biological systems has demonstrated that cellular phenotypes can be inherited across cell divisions through non-genetic mechanisms via a phenomenon termed clonal memory, where clonally related cells exhibit coordinated behaviours and fate choices that persist across multiple generations^9–13^. During reprogramming of fibroblasts to induced endodermal progenitors, early transcriptional states in sister cells predict later fate outcome^7^. Similar fate bias exists in pluripotent stem cell culture at the clonal level during formation of gastruloids, a stem cell-derived model of the gastrulating mouse embryo, where clonally related cells cultured in isolation produce gastruloids with similar phenotypes^14,15^. Together, these observations suggest that clonal memory may represent a broadly conserved mechanism by which cells encode and transmit fate information across generations, yet its role during directed differentiation of hESCs is not fully known.

The molecular basis for such fate bias remains to be fully elucidated. During hESC differentiation towards definitive endoderm (DE), signalling via the Nodal/TGFβ and WNT signalling pathways drives cells through a primitive streak-like stage towards an endoderm fate^16–19^. These pathways act in a dose-and time-dependent manner: precise levels of WNT activation initiate primitive streak formation and high levels of Activin A/Nodal signalling promote anterior primitive streak and endoderm identity. Imbalances in these signalling cues, particularly WNT, can shift differentiation outcomes away from endoderm toward mesoderm resulting in off-target populations^20–22^. Fine-tuning of signalling pathway activation does not entirely account for the variability in differentiation efficiencies^23^, raising the possibility that cell-intrinsic properties present before differentiation onset, rather than stochastic response to extrinsic signals, may underlie fate determination.

In this study, we employed clonal lineage tracing combined with single-cell multi-omics to understand the relationship between cell-cell variability in hESC differentiation towards DE. Using a lentiviral barcoding system coupled with a split-well sampling strategy during differentiation followed by scRNA-seq, we tracked individual clones across intermediate timepoints and demonstrate that clonal fate yields reproducible probabilistic outcomes. Our analysis of paired multiome snRNA-seq and snATAC-seq together with barcoding revealed that fate bias exists in hESCs prior to application of differentiation cues, where chromatin accessibility at lineage-specific cis-regulatory elements characterises this phenomenon. Specifically, we find that fate bias is associated not with enrichment of an endoderm signature in successfully differentiating clones, but rather with enrichment of accessible chromatin at mesoderm-related cis-regulatory elements in clones that produce mesodermal off-target populations. These observations have implications for stem cell differentiation efficiency and development of cell therapies by offering an explanation for how heterogeneous outcomes occur in response to the same signalling pathways.

## Results

### Clonal lineage tracing during human embryonic stem cell differentiation towards definitive endoderm

To investigate whether cell intrinsic fate bias exists during early lineage specification, we developed a clonal lineage tracing system to track individual hESC clones throughout DE differentiation. We employed a lentiviral barcoding approach, where each transduced cell receives a heritable 16-nucleotide barcode fused to an mCherry fluorescent protein driven by the strong ubiquitous promoter, EF1ɑ^24^, randomly integrated into its genome. This enabled tracking of clonal relationships through cell division and differentiation **(Fig. 1A)**. To validate barcode complexity and determine barcode collision rates within our experimental setup, we transduced 4×10^3^ hESCs maintained in defined serum-free pluripotent stem cell medium (Nutristem) across two parallel replicate transductions. After 7 days in culture, these cells expressed high levels of mCherry by fluorescence microscopy **(Suppl. 1A)**. mCherry-positive hESCs were isolated from each replicate by fluorescence-activated cell sorting (FACS) **(Suppl. 1B)** followed by scRNA-seq **(Suppl. 1C)**. Comparison of barcodes between replicates revealed minimal overlap (<1%; 14 shared barcodes among 1,542 clone calls across the two replicate populations), confirming negligible barcode collision and validating our clone calling pipeline **(Suppl. 1D)**^7,25^.

**Figure 1:**
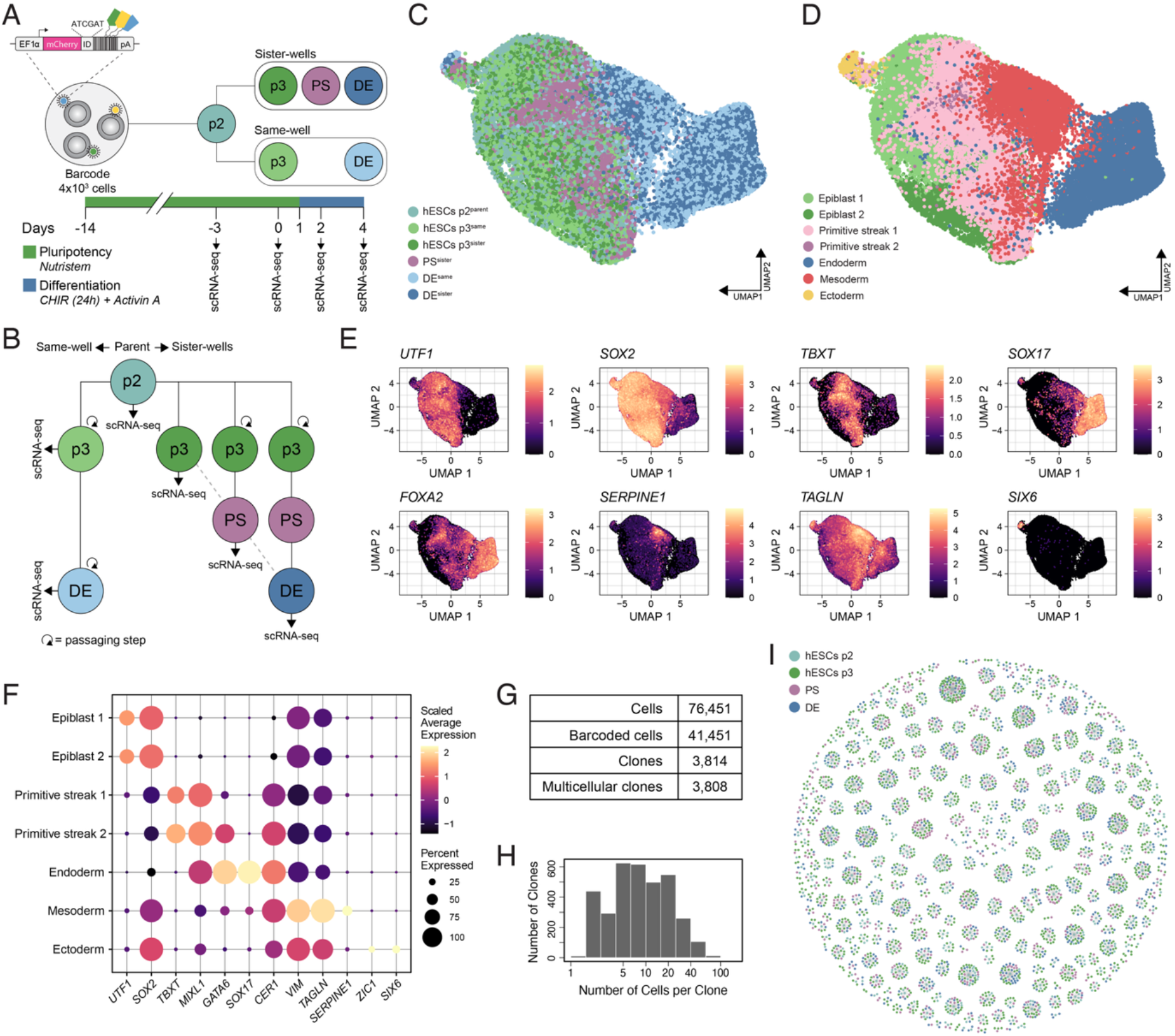
Clonal lineage tracing during human embryonic stem cell differentiation towards definitive endoderm. (A) Schematic of experimental design. 4×10^3^ hESCs were transduced with the lentiviral barcode library, expanded and differentiated towards definitive endoderm (DE) using 3µM CHIR99021 and 100ng/mL Activin A. Timeline indicates culture conditions and scRNA-seq sampling timepoints. (B) Split-well sampling strategy. At hESCs p2, cultures were split into four wells. Following passage to hESCs p3, one well was sampled before differentiation and again at DE (hESCs p3, DE; same-well strategy), while the remaining wells were sampled independently at hESCs p3, primitive streak (PS), or DE (sister-well strategy), enabling temporal sampling while testing clonal reproducibility across independently cultured replicates. (C, D) UMAP dimensional embedding of RPCA-integrated scRNA-seq coloured by (C) timepoint and sampling strategy, and (D) cell type annotation based on unsupervised clustering. (E) UMAP dimensional embedding showing log-normalised expression of indicated markers. (F) Dot plot graph showing scaled normalized average expression of indicated markers across cell types. Dot size represents percentage of expression; colour intensity represents scaled normalised average expression. (G) Numbers of cells and clones sampled in the dataset. (H) Distribution of clone sizes (cells per clone). Median = 8 cells; mean = 22 cells; range 1-101 cells. (I) Force-directed graph visualisation of clonal relationships. Each circle represents on cell coloured by timepoint; cells from the same clone are connected, showing clonal structure across developmental stages (*n* = 381 clones; 10% of total number of clones).

In order to maximise temporal resolution during hESC to DE differentiation, while avoiding technical confounds from repeated sampling, we devised a split-well sampling strategy **(Fig. 1A, B)**. Parental hESCs (4×10^3^ cells) were transduced with the barcode library and expanded to passage 2 (hESC p2, Day -14 to -3), reaching ∼2×10^6^ cells through roughly 9 cell doublings **(Suppl. 1E)**. This expansion ensured that each barcode founder cell gave rise to a multi-cell clone. The culture was then split into four replicate wells, such that each clone was represented in multiple wells. These wells were subsequently sampled at different timepoints, where one well was resampled at multiple timepoints (same-well strategy: hESC at Day 0 and DE at Day 4), while the remaining three were split into sister wells, each sampled only once at a given timepoint (sister-well strategy: hESC at Day 0, primitive streak (PS) at Day 2, and DE at Day 4). This dual approach allowed us to sample at an additional timepoint (PS) during DE differentiation, while avoiding introducing biological variations from repeated sampling of the same well. More importantly, because sister wells share clonal composition by design, this strategy enabled testing whether clonal behaviours are reproducible across spatially segregated but genetically identical replicate cultures. This provides the inferential foundation for distinguishing heritable cell-intrinsic fate bias from stochastic variation arising during differentiation or sampling. Following the split at hESC p2 (day -3), cells were passaged once more at hESC p3 (Day 0) and cultured in Nutristem medium for an additional 24h, after which DE differentiation was induced by replacing medium with RPMI 1640 supplemented with Activin A and CHIR99021 (CHIR; Day 1 to 2). Following a 24h pulse with CHIR to drive primitive streak formation (Day 1 to 2)18,26, CHIR was removed and the culture was maintained in Activin A alone from Day 2 until Day 4.

We performed scRNA-seq across 4 timepoints for a total of 6 samples between same and sister wells: hESC p2 (hESC p2^parental^), hESC p3 (hESC p3^same^ and hESC p3^sister^), primitive streak (PS^sister^, Day 2) and definitive endoderm (DE^same^ and DE^sister^, Day 4). During quality control and filtering, we observed a subpopulation within the female hESC culture with low *XIST* expression characteristic of X-chromosome inactivation erosion (XCIE) **(Suppl. 1F, G)**. As XCIE represents a culture-adapted state that can influence differentiation trajectories^27^, we performed hierarchical sub-clustering of pluripotent populations to identify and exclude these cells **(Suppl. 1H, I)**. Clonal tracking confirmed that XCIE clones exhibited distinct lineage trajectories compared to *XIST*-expressing clones **(Suppl. 1J, K)**, validating this filtering strategy. All subsequent analyses were performed on this refined dataset. Unsupervised clustering revealed distinct transcriptional states corresponding to the expected developmental stages, visualised by UMAP embedding **(Fig. 1C, D)**. We identified two epiblast-like populations (Epiblast 1, Epiblast 2) spanning hESC p2 and p3 timepoints, two primitive streak populations (Primitive streak 1, Primitive streak 2), and three representative germ layer populations (Endoderm, Mesoderm and Ectoderm). Expression of canonical lineage markers confirmed these representative cell type annotations: *UTF1* and *SOX2* marked pluripotent cells; *TBXT* marked primitive streak; *SOX17* and *FOXA2* marked endoderm; *SERPINE1* and *TAGLN* marked mesoderm; and *SIX6* marked ectoderm **(Fig. 1E, F)**. We recovered 76,451 cells, of which 41,451 (54.2%) contained identifiable barcodes representing 3,814 unique clones **(Fig. 1G)** with clone sizes ranging from 1 to 101 cells **(Fig. 1H)**. Force-directed graph visualisation of a representative subset of clones in the dataset (10%) revealed extensive clonal structure **(Fig. 1I)**. Despite directed differentiation of hESCs towards DE through stimulation of TGFβ and WNT-related signalling, we observed a substantial mesoderm off-target population (∼39% of cells at Day 4) **(Fig. 1D; Suppl. 1L)**, providing an opportunity to investigate whether this heterogeneity reflects stochastic fate decisions or cell-intrinsic biases present prior to onset of differentiation using our clonally-resolved dataset.

### Clonally related cells exhibit reproducible fate bias across independent split-well culture replicates

To determine whether clonally related cells exhibit coordinated fate decisions, we first assessed transcriptional similarity between cell pairs *within* versus *between* clones at the earliest sampled timepoint 11 days post-transduction (hESCs p2). Cell pairs from the same clone showed significantly higher transcriptional correlation compared to randomly sampled cell pairs from different clones (Cohen’s d = 3.51, *p* < 0.0001) **(Fig. 2A)**. This indicates that clonal identity is associated with shared molecular features that are inherited stably across multiple cell divisions and spatial segregation of clonal sisters during culture expansion, with within-clone similarity persisting to at least hESCs p3 **(Fig. 2B)**. We ensured robust statistical analysis of clonal data by applying a filtering strategy requiring detection at both undifferentiated stages (hESC p2, hESC p3), ≥4 cells per clone, detection at ≥2 timepoints, continuous temporal coverage without gaps indicative of resurrecting clones and ≥2 cells per detected timepoint **(Suppl. 2A)**. To distinguish clones that successfully transition through differentiation from those that fail to expand, we categorised clones as “winner” (detected at all timepoints with sustained growth) or “loser” (lost during differentiation) clones **(Fig. 2C; Suppl. 2A)**. Clone distributions were robust to the choice of Jaccard similarity threshold used for clone calling, with <2% variation in clone counts across thresholds of 0.6-0.8 **(Suppl. 2B)**. Winner clones substantially outnumbered loser clones (1,635 versus 397 clones) **(Fig. 2D; Suppl. 2B)**, containing overall larger cell numbers per clone **(Suppl. 2C)**. However, differential gene expression analysis between winner and loser clones at hESCs p3 revealed minimal transcriptional differences **(Suppl. 2D, E)**. These differences in clone sizes may be due to either low sampling rates of loser clones or neutral drift dynamics with stochastic clone loss accompanied by compensatory expansion of surviving clones^28^. This indicates that loser clones are not transcriptionally distinct from winner clones at the pluripotent stage. Subsequent fate-bias analyses therefore focused on winner clones, since loser clones do not persist to the differentiation timepoints where fate is assayed. Winner clones underwent further filtering for barcodes shared between same-well and sister-wells at hESCs p3 and DE **(Fig. 2E, F)**, yielding a total of 981 clones for analysis. Clones that were removed due to not overlapping across same-well and sister-well replicates were found to have significant lower cell numbers, consistent with stochastic sampling dropout **(Suppl. 2F)**.

**Figure 2:**
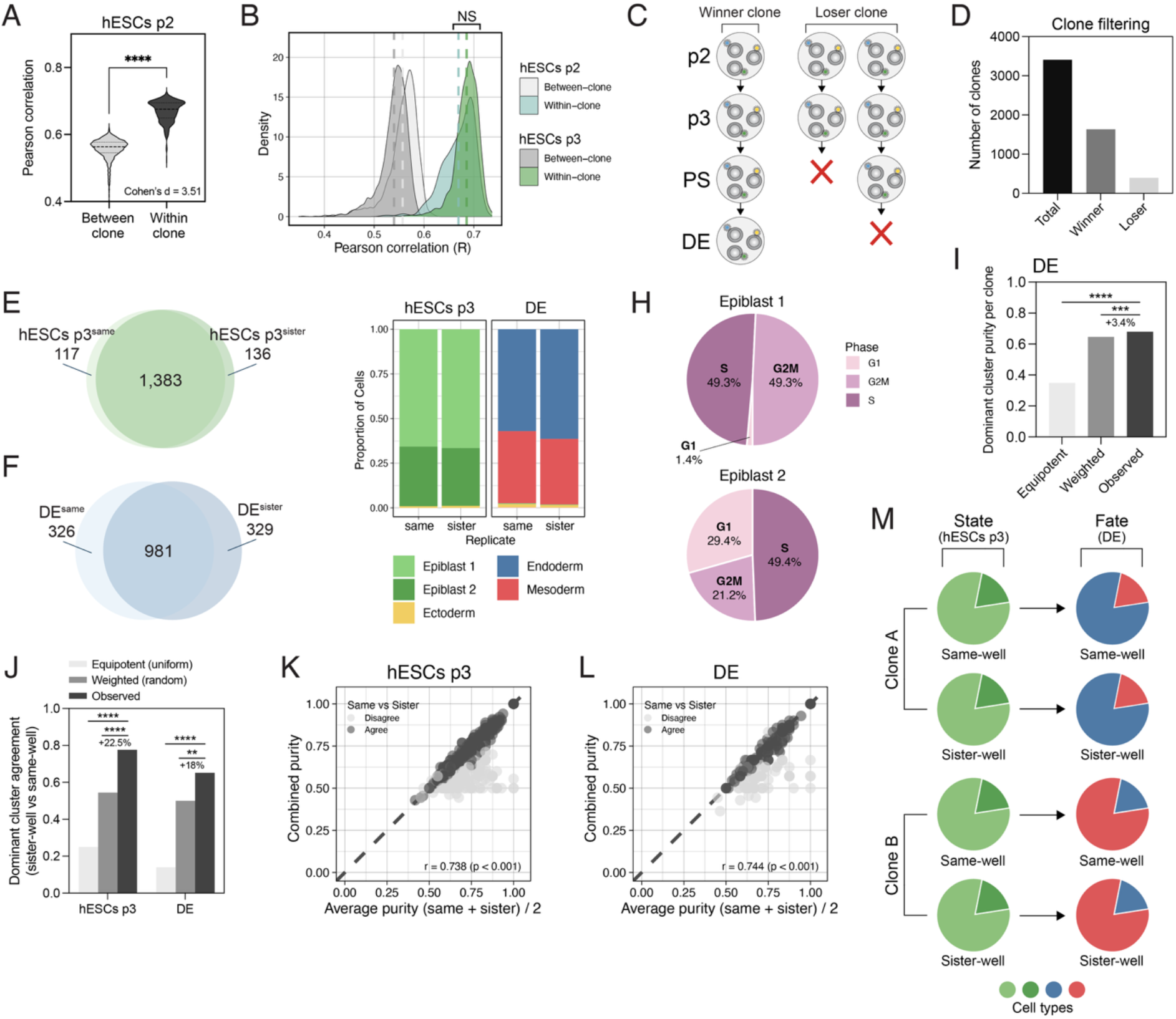
Clonally-related cells exhibit reproducible fate bias across independent replicates. (A) Violin plot of transcriptional similarity (Pearson correlation) between cell pairs at hESCs p2. Within-clone pairs show significantly higher correlation that random between-clone pairs (Cohen’s d = 3.51, *p* < 0.0001, Wilcoxon test). (B) Distribution of Pearson correlations for within-clone vs between-clone pairs at hESCs p2 and p3. Increased within-clone similarity persists through passage (NS, not significant between timepoints). (C) Schematic defining “winner” clones (detected at all timepoints) vs “loser” clones (lost during differentiation). Red X indicates clone dropout. (D) Clone filtering summary. Of 3,814 total clones, 1,635 were winners and 397 clones were losers. The remaining 1,782 clones were filtered out due to not passing minimum clone thresholds defined in Suppl. 2A. (E, F) Venn diagrams showing clone overlap between same-well and sister-well replicates at (E) hESCs p3 and (F) DE. (G) Stacked bar plots comparing cluster composition across same-well and sister-well replicated at hESCs p3 and DE (hESCs p3: Cramér’s V < 0.048, p > 0.05; DE: Cramér’s V < 0.05, *p* < 0.001). (H) Pie charts showing cell cycle phase composition of Epiblast 1 and Epiblast 2 clusters at hESCs p3. Colour represents cell cycle phases (G1, S, G2M). (I) Bar plot showing population-level analysis of cluster purity within clone. Observed purity (proportion of cells in dominant cluster) is significantly higher than expected from weighted random sampling of the cluster distribution with small effect size (+3.4 percentage points, *p* < 0.001). (J) Bar plot of dominant cluster agreement between same-well and sister-well replicates. Observed agreement is significantly higher than weighted random expectation at both hESCs p3 (77% vs 54.5%, *p* < 0.0001) and DE (68.7% vs 50%, *p* < 0.01, Chi-square test). (K, L) Scatter plots comparing clone purity calculated after pooling cells across same-well and sister-well replicates with the mean purity calculated independently in each replicate at (K) hESCs p3 and (L) DE. The two approaches were strongly correlated at both stages (hESCs p3: r = 0.738, *p* < 0.001; DE: r = 0.744, p < 0.001). Points coloured by purity difference; dashed line indicates identity. Agree and disagree classifications shown in legend. (M) Schematic summary showing that clonally-related cells reproducibly have the similar dominant cluster fate agreement at both hESCs (state) and DE (fate) samples, validating pooling of clones from both sampling strategies for downstream analysis.

We assessed the reproducibility of cell type composition across independently cultured replicates by first comparing same-well and sister-wells at the cluster level **(Fig. 2G)**. The two epiblast-like clusters in hESCs p3 were primarily distinguished by cell cycle state, with cells in Epiblast 1 actively proliferating, and Epiblast 2 containing a sizeable population exiting cell cycle **(Fig. 2H)**. At DE, cells differentiated towards either endoderm or mesoderm **(Fig. 2G)**. Across both sampling timepoints, a small subpopulation (<5%) of ectoderm-like cells was consistently observed. When comparing proportions of cells in each cluster per timepoint between same-well and sister-wells, we observed negligible effect sizes (hESCs p3: Cramér’s V = 0.048, *p* > 0.05; DE: Cramér’s V = 0.05, *p* < 0.001), indicating that population-level differentiation outcomes are reproducible between replicates.

We then leveraged the clone-level resolution of our data and quantified clonal coherence as a purity metric defined by the fraction of cells within each clone assigned to the same dominant cluster at a given timepoint **(Suppl. 2A; Step 3-4)**. When comparing observed purity values across all clones at the DE stage to an equipotent null model, we found that clones exhibited significantly higher purity than random expectation (observed: 0.68 versus random weighted: 0.646, *p* < 0.001, *n* = 495 clones) **(Fig. 2I)**. While this indicated that clonally related cells display coordinated fate choices during differentiation, the magnitude of this effect was minimal (+3.4%), suggesting substantial clone-clone variability that could arise either from true biological heterogeneity or stochastic sampling noise. To distinguish between these two possibilities, we took advantage of our split-well experimental design **(Fig. 1B)** and adopted a categorical approach that asked whether fate preferences are reproducible across genetically identical but spatially segregated replicates based on dominant cluster agreement. We observed dominant cluster agreement between hESCs p3^same^ and hESCs p3^sister^ clones (0.77 versus 0.54 random weighted, Chi-square *p* < 0.0001, *n* = 343 clones) **(Fig. 2J)**. As hESC p3 cells primarily cluster based on cell cycle **(Fig. 2H)**, this indicates that clonally related hESCs exhibit coordinated cell cycle dynamics that are maintained even across spatially separated culture wells, consistent with previous reports in ESCs^29–31^. We observed a similar pattern of agreement between dominant fates across DE^same^ and DE^sister^ clones (0.69 versus 0.50 random weighted, Chi-square *p* < 0.01, *n* = 67 clones) **(Fig. 2J)**. This high degree of reproducibility demonstrates that clonally related cells maintain coordinated fate choices during differentiation. This replicate-based strategy provides orthogonal evidence for cell-intrinsic fate bias, whereby clones exhibiting consistent dominant fate identities across independently cultured replicates support heritable fate preference rather than arising from sampling noise.

To quantitatively assess the agreement between sampling strategies across both stages, we calculated purity correlations **(Fig. 2K, L)**. Purity values computed using two different pooling strategies (per-replicate then averaged vs cells pooled across replicates) were highly correlated at both stages (hESCs p3: *r* = 0.738, *p* < 0.001; DE: *r* = 0.744, *p* < 0.001). Bland-Altman analysis confirmed minimal systematic bias between sampling methods at both timepoints (mean difference ≈ 0; limits of agreement; ±1.96 SD) **(Suppl. 2G)**. This further confirms that same-well and sister-well strategies yield equivalent measurements of clonal fate bias and validates pooling clones from both sampling approaches for downstream analysis. We then asked the extent to which clone size affects clonal coherence metrics by calculating Euclidean distances between cells within each clone **(Suppl. 2H)**. Larger clones showed decreased Euclidean distances across cells at both hESCs p3 and DE, with increased sampling likely reducing noise that would otherwise be present in smaller clones. We additionally confirmed that the observed clone-level purity is not positively inflated by clone size does not correlate with differences in clone size at both hESCs p3 (Spearman *r* = −0.09, *p* < 0.01) and DE (Spearman *r* = −0.16, *p* < 0.001) stages **(Suppl. 2I, J)**. These findings underscore the importance of adequate clone sizes for accurate lineage tracing analysis. Together, these analyses confirm that same-well and sister-well strategies yield equivalent measurements of clonal fate bias and related clones were pooled for all further analyses **(Fig. 2M)**.

### Fate bias towards endoderm or mesoderm lineages is not captured by early transcriptomic signatures

Having established that clonally related cells exhibit reproducible fate biases during hESC differentiation, we next sought to characterise the spectrum and directionality of these biases during germ layer specification. As endoderm and mesoderm constitute the dominant cell fates at the DE stage (59% and 39% of cells, respectively) **(Fig. 2G)**, with ectoderm representing a minor persistent off-target population (<2%) **(Suppl. 3A-C)**, we focused our analysis on endoderm versus mesoderm fate allocation.

Hierarchical clustering of clones based on their endoderm or mesoderm composition revealed reciprocal lineage allocation at a clonal level **(Fig. 3A)**. Clones segregated into two major groups: those strongly biased towards endoderm (high endoderm fraction, low mesoderm fraction) and those strongly biased toward mesoderm (high mesoderm fraction, low endoderm fraction). This pattern indicates that most clones are biased towards a dominant fate rather than bipotential. We then calculated a fate bias score for each clone to discern lineage preference at the clonal level, defined as the proportion of cells adopting an endoderm fate minus the proportion adopting a mesoderm fate at the DE stage, to understand how our clones might broadly fit into these models: a bimodal distribution capturing fate bias or a unimodal distribution capturing bipotency **(Fig. 3B)**. The distribution of fate bias scores across all clones revealed a bimodal pattern, where most clones were either dominantly (>70%) endoderm or mesoderm and less commonly a balance between both. Endoderm-biased and mesoderm-biased clones pass through a shared primitive streak intermediate before diverging to their respective germ layer fates at the DE stage **(Fig. 3C)**.

**Figure 3:**
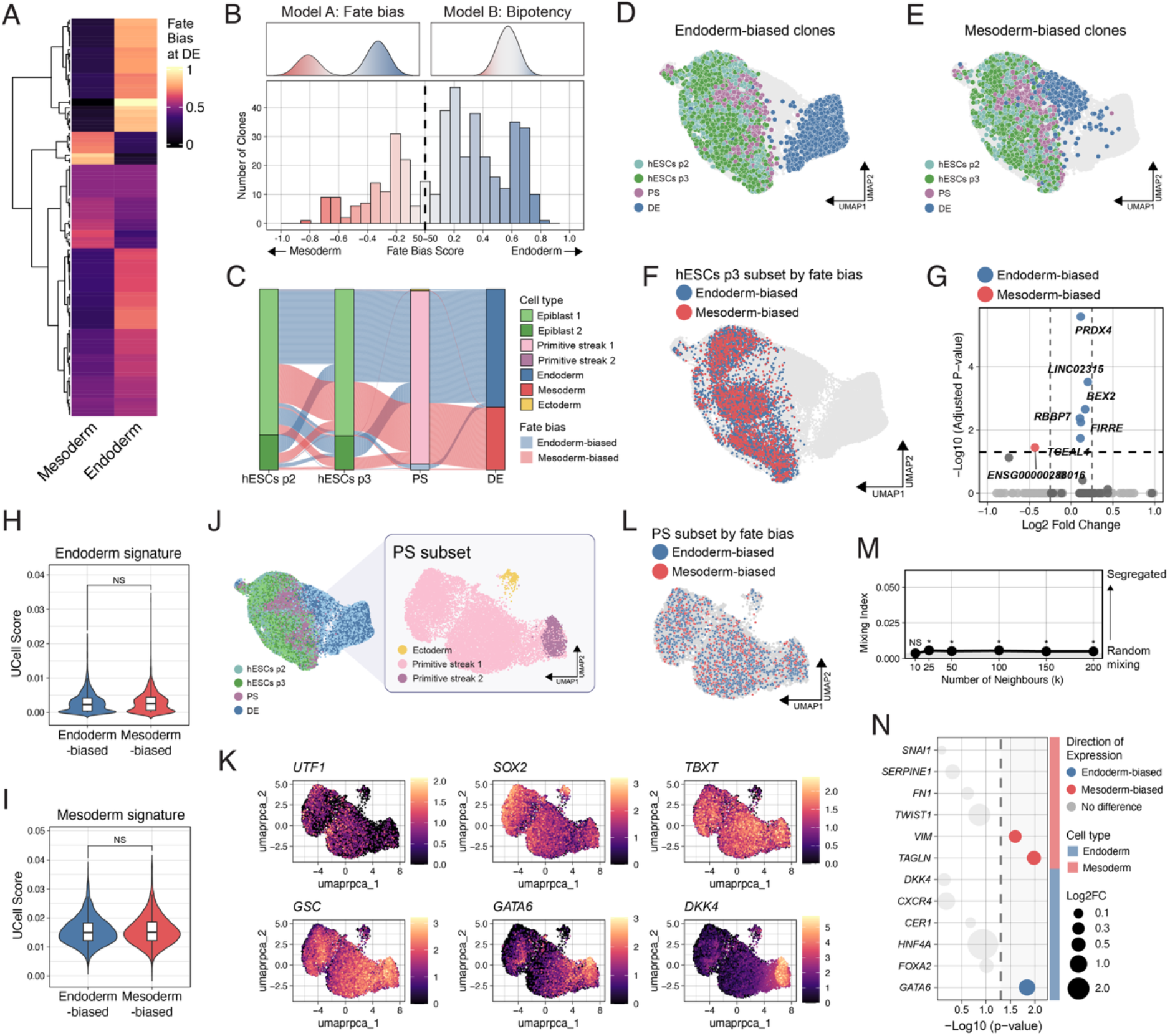
Clones exhibit fate bias towards either endoderm or mesoderm cell types not explained by transcriptomic signatures at earlier timepoints. (A) Heatmap of hierarchical clustering of clones based on endoderm or mesoderm composition at DE. Each row represents one clone. Colour intensity indicates fate bias (purple = mesoderm-biased, yellow = endoderm-biased). (B) Histogram of fate bias scores across all clones. Fate bias score = (proportion of cells in endoderm) – (proportion in mesoderm), ranging from −1 (fully mesoderm clones) to +1 (fully endoderm clones). Dashed line indicates balanced bipotent clones. (C) Sankey plot showing clone trajectories for endoderm-biased and mesoderm-biased clones across all sampled developmental stages. Streams are coloured by fate bias and strata are coloured by annotated cell type. (D, E) UMAP dimensional embedding visualising clone trajectory of all (D) endoderm-biased clones and (E) mesoderm-biased clones. (F) UMAP dimensional embedding of all endoderm-biased and mesoderm-biased clones at hESCs p3 overlayed. (G) Volcano plot of differential gene expression between endoderm-biased and mesoderm-biased clones at hESCs p3. (H, I) Violin plots showing UCell signature scores for (H) endoderm and (I) mesoderm gene signatures in endoderm-biased and mesoderm-biased clones at hESCs p3. (J) UMAP embedding of (left) full dataset and (right) the PS subset re-embedded for visualisation. Cluster annotations (Primitive streak 1, Primitive streak 2, Ectoderm) were retained from the full dataset analysis. (K) UMAP dimensional embedding showing log-normalised expression of indicated markers. (L) UMAP dimensional embedding of all endoderm-biased and mesoderm-biased clones at PS overlayed. (M) Mixing index quantifying spatial segregation of fate-biased clones within the PS subset across varying neighbourhood sizes (k=10-200 nearest neighbours). Values near 0 indicate random mixing; positive values indicate segregation. (N) Bubble plot showing expression of canonical endoderm (top) and mesoderm (bottom) markers between endoderm-biased and mesoderm-biased clones at the PS stage. Bubble colour indicates direction of expression (upregulated in endoderm-biased or mesoderm-biased clones); bubble size indicates magnitude of expression; dashed line indicates *p* = 0.05.

To identify molecular features associated with fate bias, we first subset endoderm-biased and mesoderm-biased clones based on the stringent criteria that all cells within a clone give rise to either endoderm or mesoderm at DE **(Fig. 3D, E)**. We note that this threshold is an analytical choice to enrich for clones with clear directional bias and does not imply that fate allocation is biologically binary, where the continuous distribution of fate biased scores **(Fig. 3B)** reflects a spectrum of outcomes. We then performed differential gene expression analysis between endoderm-biased and mesoderm-biased clones at hESCs p3, 14 days post-transduction and before differentiation cues were applied **(Fig. 3F, G)**. Here, we observed few differentially expressed genes (DEGs), where those that were found to be significant had negligible effect sizes (log2 fold change < 0.25, with the exception of TCEAL4). We then tested whether fate-biased clones differed in their overall lineage-specific transcriptional programmes rather than at the level of individual genes by calculating signature scores^32^ for endoderm and mesoderm gene sets. However, neither the endoderm **(Fig. 3H)** nor the mesoderm **(Fig. 3I)** signature scores showed significant differences between endoderm-biased and mesoderm-biased clones at hESCs p3, indicating that overall, our analyses could not resolve fate bias transcriptionally by scRNA-seq at this stage.

### Subtle transcriptional changes emerge during PS stage reflective of future fate bias

As migration through the primitive streak during gastrulation results in emergence of cells towards different germ layers, we next examined cell dynamics at the critical PS stage corresponding to embryonic gastrulation where cells migrate to specific germ layers **(Fig. 3J)**. Expression of candidate lineage markers demonstrated that cells within the *TBXT*-expressing PS timepoint separated temporally through primitive streak development, with cells located on the left side of the UMAP representing an earlier primitive streak cell type, based on residual expression of pluripotency markers such as *UTF1* and *SOX2*, while cells located on the right side expressed markers of the posterior primitive streak such as *DKK4* **(Fig. 3K)**. These expression patterns demonstrate that cells within the PS timepoint separate primarily along a temporal axis representing progressive primitive streak maturation. Endoderm-biased and mesoderm-biased clones overlayed onto the PS subset showed extensive spatial overlap **(Fig. 3L)**, indicating no preferential segregation into distinct germ layer substates. This was quantified by calculating mixing scores across varying neighbourhood sizes (k = 10-200 nearest neighbours) **(Fig. 3M)**, where scores remained near zero with negligible effect sizes despite significant segregation at larger neighbourhood sizes, confirming that most endoderm-biased and mesoderm-biased clones are randomly distributed rather than spatially segregated at the PS stage. Despite this general spatial mixing of clones, when performing differential expression analysis for lineage-specific markers between endoderm-biased and mesoderm-biased clones within the PS subset, we observed enrichment of the endoderm marker *GATA6* expression in endoderm-biased clones and enrichment of the mesodermal mesenchyme markers *VIM* and *TAGLN* in mesoderm-biased clones **(Fig. 3N)**. This suggests that already at the PS stage, subtle lineage-specific transcriptional differences begin to emerge between endoderm-biased and mesoderm-biased clones. These results indicate that if transcriptional differences between fate-biased clones exist in hESCs prior to the onset of differentiation, then they are below the detection threshold of our scRNA-seq approach, pointing instead towards an alternative regulatory layer as the molecular basis for heritable fate bias in hESCs.

### Chromatin accessibility at mesoderm-specific regions is associated with differentiation outcome in response to WNT and TGFβ signalling

Epigenetic modifications are heritable states that can be propagated across many generations and demonstrated as a route for conveying memory and fate potential in various biological systems^33–36^. We asked whether fate bias in hESCs prior to differentiation may be encoded across the epigenome by assessing chromatin accessibility at lineage-specific cis-regulatory elements^33^. We generated a multiome by performing paired single-nucleus RNA-seq (snRNA-seq) and single-nucleus Assay for Transposase Accessible-Chromatin-seq (snATAC-seq) on barcoded nuclei sampled at hESCs p3 (Day 0), PS (Day 2) and DE (Day 4) **(Fig. 4A)**. This approach enabled simultaneous profiling of the transcriptome and chromatin accessibility within the same nuclei, allowing direct integration of gene expression with regulatory landscapes at the clonal level. Following quality control and filtering, this dataset yielded 12,741 nuclei, of which 4,323 contained identifiable barcodes representing 1,333 unique clones with clone sizes ranging from 1-40 cells per clone **(Fig. 4B, C)**. Cell type annotations from unsupervised clustering of the snRNA-seq dataset visualised by UMAP embedding **(Fig. 4D; Suppl. 4A-C)** broadly recapitulated those identified in the previous scRNA-seq dataset **(Fig. 1D)**, with hESCs segregating into epiblast (Epiblast 1 and Epiblast 2), primitive streak, endoderm, mesoderm and ectoderm (Ectoderm 1 and Ectoderm 2) populations. While unsupervised clustering based on chromatin accessibility within the snATAC-seq dataset was mostly concordant with the transcriptome, the primitive streak snRNA-seq cluster was split into two snATAC-seq clusters (Primitive streak 1_atac and Primitive streak 2_atac) **(Fig. 4D; Suppl. 4D)**. Genomic annotation of differentially accessible peaks (DAPs) across all cell types showed that the majority of peaks localised to distal intergenic or intronic regions, consistent with enhancer-mediated regulation **(Suppl. 4E)**.

**Figure 4:**
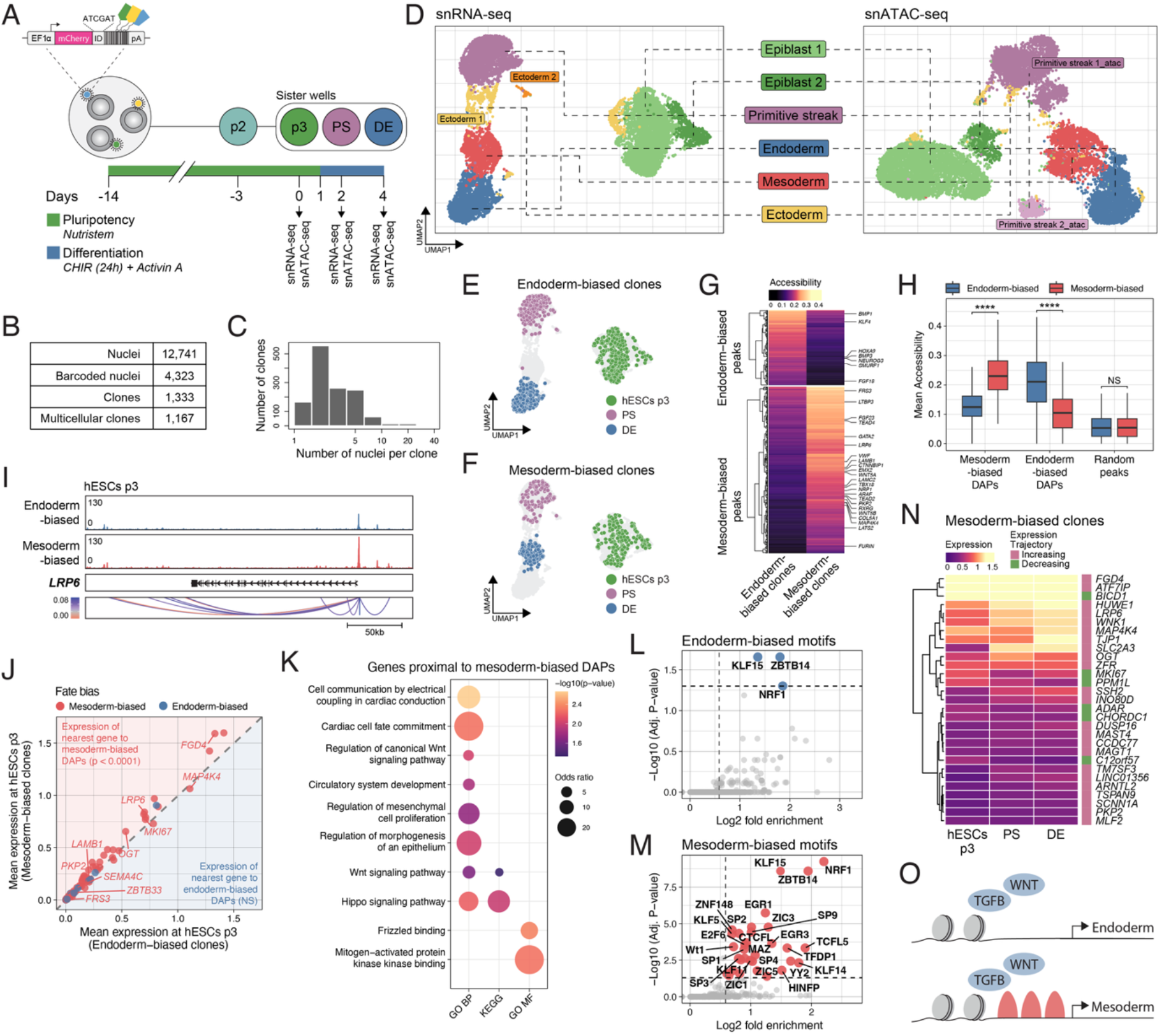
Chromatin accessibility at mesoderm-specific regions biases differentiation outcome in response to WNT and TGFβ signalling. (A) Schematic of paired snRNA-seq/snATAC-seq (multiome) experimental design. 4×10^3^ hESCs barcoded with the lentiviral barcode library, expanded and differentiated towards definitive endoderm (DE) using 3µM CHIR99021 and 100ng/mL Activin A. Timeline indicates culture conditions and snRNA-seq sampling timepoints. (B) Numbers of nuclei and clones sampled in the dataset. (C) Distribution of clone sizes in the multiome dataset. (D) UMAP dimensional embeddings of snRNA-seq (left) and snATAC-seq (right) datasets coloured by cell type annotation based on unsupervised clustering for RNA and ATAC assays, respectively. Dashed lines highlight corresponding populations between RNA and ATAC modalities. (E, F) UMAP dimensional embedding visualising clone trajectory of all (E) endoderm-biased clones and (F) mesoderm-biased clones. (G) Heatmap showing mean chromatin accessibility at endoderm-biased peaks (top) and mesoderm-biased peaks (bottom) across all nuclei in endoderm-biased and mesoderm-biased clones. Colour indicates average accessibility score. (H) Box plot comparing mean chromatin accessibility between mesoderm-biased (left pair) and endoderm-biased (middle pair) DAPs in endoderm-biased vs mesoderm-biased clones and random genomic peaks as a control (right pair) at hESCs p3. (I) Coverage plot showing chromatin accessibility at the LRP6 locus in endoderm-biased clones (top) versus mesoderm-biased clones at hESCs p3. Black boxes indicate accessible peak regions; arcs indicate peak-to-gene links; scale bar indicates genomic distance. (J) Scatter plot comparing mean expression of the nearest gene to mesoderm-biased DAPs (red, *p* < 0.0001) and endoderm-biased DAPs (blue, NS) between mesoderm-biased clones (y-axis) and endoderm-biased clones (x-axis) at hESCs p3. Points above the diagonal line indicate higher expression in mesoderm-biased clones. (K) Gene ontology (GO Biological Process (BP), KEGG, GO Molecular Function (MF)) enrichment analysis of genes nearest to mesoderm-biased DAPs. Dot size represents odds ratio; colour intensity represents −log10(p-value). (L, M) Scatter plot of transcription factor motif enrichment in (L) endoderm-biased DAPs and (M) mesoderm-biased DAPs. (N) Heatmap showing expression trajectories across developmental stages of genes proximal to mesoderm-biased DAPs in mesoderm-biased clones. Rows represent individual DAP-proximal genes hierarchically clustered by expression pattern. Rows represent individual genes hierarchically clustered by expression pattern. Colour bar indicates expression level (magenta = increasing; green = decreasing). (O) Schematic showing proposed epigenetic fate priming mechanism. Prior to differentiation (hESCs p3), clones exhibit differential chromatin accessibility at lineage-specific regulatory regions despite similar transcriptional states. In response to TGFβ and WNT signalling, endoderm-biased clones (top) respond appropriately to differentiation cues, while mesoderm-biased clones (bottom) harbour mesoderm-specific accessibility resulting in preferential activation of mesoderm programmes.

Using the same clone filtering and fate bias scoring strategy as for the scRNA-seq dataset **(Suppl. 2A)**, we identified endoderm-biased (*n* = 390 clones) and mesoderm-biased clones (*n* = 211 clones) in the multiome dataset. Visualisation of these fate-biased clones across developmental stages confirmed their distinct trajectories **(Fig. 4E, F)**. Differential expression analysis between endoderm-biased and mesoderm-biased clones at hESCs p3 did not find any DEGs **(Suppl. 4F)**, recapitulating the observations made in the scRNA-seq dataset **(Fig. 3F-I)**. To identify chromatin features associated with fate bias, we performed differential chromatin accessibility analysis between endoderm-biased clones and mesoderm-biased clones at hESCs p3, resulting in 181 endoderm-specific DAPs and 402 mesoderm-specific DAPs **(Fig. 4G)**. This asymmetry in the total number of DAPs and in the magnitude of accessibility differences may reflect more extensive epigenetic changes in mesoderm-biased clones. We first confirmed that observed differences reflected localised rather than global chromatin changes, where total chromatin accessibility **(Suppl. 4G)** and transcription start site (TSS) enrichment scores **(Suppl. 4H)** showed no significant differences between fate-biased clones. To validate that DAPs represent fate-specific regulatory elements, we compared mean chromatin accessibility at mesoderm-biased DAPs, endoderm-biased DAPs, and randomly sampled genomic peaks between fate-biased clones **(Fig. 4H, I; Suppl. 4I)**. We found that mesoderm-biased clones showed significantly higher accessibility at mesoderm-specific DAPs compared to endoderm-biased clones (*p* < 0.0001), and endoderm-biased clones showed significantly higher accessibility at endoderm-specific DAPs (*p* < 0.0001), while randomly sampled peaks showed no significant difference between fate-biased clones, confirming that DAP signal is locus-specific rather than reflecting global chromatin variation between clone groups.

Chromatin accessibility at cis-regulatory elements can reflect distinct regulatory states, including poised regulatory elements primed for future activation and active transcriptional regulation of already-expressed genes^37,38^. To discern between these two possibilities at fate-biased DAPs, we first examined the relationship between differential chromatin accessibility and expression of DAP-proximal genes at hESCs p3 **(Suppl. 4J)**. When comparing mean expression of nearest genes to fate-biased DAPs for endoderm-biased and mesoderm-biased clones against randomly selected genes, we found that only nearest genes to mesoderm-biased DAPs showed significantly higher expression (*p* < 0.0001) **(Suppl. 4K)**. Direct comparison of DAP-proximal genes in endoderm-biased clones versus mesoderm-biased clones similarly showed higher expression of genes proximal to mesoderm-biased DAPs compared to endoderm-biased clones (*p* < 0.0001) **(Fig. 4J)**. Although nearest-gene annotation may not capture the true regulatory targets of distal peaks, these analyses demonstrate that differential chromatin accessibility in mesoderm-biased clones, not endoderm-biased clones, is accompanied by subtle but consistent differences in proximal gene expression at hESCs p3 that fall below the threshold of individual DEG detection, potentially predisposing these towards distinct responses upon receiving differentiation cues.

We next investigated whether mesoderm-biased DAPs localise near genes involved in mesoderm development and signalling pathways, as we had previously identified the promoter for the WNT co-receptor *LRP6* as differentially accessible in mesoderm-biased clones **(Fig. 4I)**. Given that the DE differentiation protocol employs the GSK3β inhibitor, CHIR, that activates WNT signalling by preventing degradation of β-catenin, pre-existing accessibility at WNT pathway-related genes, such as *LRP6*, could lead to hypersensitivity to stimulation, resulting in preferential mesoderm over endoderm fate specification^20–22^. Pathway enrichment analysis of nearest genes to mesoderm-biased DAPs revealed significant enrichment of WNT signalling, cardiac cell fate commitment, and regulation of mesenchymal cell proliferation **(Fig. 4K)**. To systematically identify transcription factor (TF) binding motifs enriched near fate-biased DAPs, we performed motif enrichment analysis **(Fig. 4L, M)**. Endoderm-biased DAPs showed enrichment for only 3 TF motifs, all of which were shared with mesoderm-biased DAPs: KLF15, ZBTB14 and NRF1 **(Fig. 4L)**. These may be broadly expressed TFs associated with general chromatin organisation rather than lineage-specific developmental programmes. By contrast, mesoderm-biased DAPs showed enrichment for an additional >30 TF motifs **(Fig. 4M)**. This is consistent with mesoderm-biased clones harbouring pre-accessible regulatory sites that facilitate preferential activation of mesoderm programmes in response to signalling.

To link chromatin accessibility with subsequent gene expression changes during differentiation, we examined the expression trajectories of nearest genes proximal to mesoderm-biased DAPs across developmental stages in mesoderm-biased clones **(Fig. 4N; Suppl. 4L-N)**. Hierarchical clustering revealed distinct expression patterns, with a majority of regulated genes increasing in expression over developmental time **(Fig. 4N)**. When performing a similar analysis for expression trajectories of nearest genes to endoderm-biased DAPs, we observed only subtle expression differences across time **(Suppl. 4N).** When looking at the perdurance of accessible chromatin sites across fate-biased clones between hESCs p3 and PS stages, we observed persistence of approximately 11% of mesoderm-biased DAPs from hESC p3 to PS, compared with approximately 5% of endoderm-biased DAPs **(Suppl. 4L, M)**. Together, these temporal analyses leverage the clonal resolution of our data to show that differences in chromatin accessibility at the pluripotent stage prefigure gene expression changes during differentiation. Overall, these results support an asymmetric fate priming model in hESCs whereby pre-existing chromatin accessibility at mesoderm-specific regulatory regions, not endoderm, is associated with differentiation outcome in response to WNT and TGFβ signalling **(Fig. 4O)**.

## Discussion

In this study, integration of clonal lineage tracing with single-cell multiomics reveals that heterogeneous outcomes during directed differentiation of hESCs are associated with chromatin features in undifferentiated cells established long before differentiation signals are applied. Central to our approach is a split-well sampling design that allowed clonal fate preferences to be measured across spatially segregated replicates, providing the inferential foundation for the heritability of clone-level fate bias. Coupled with scRNA-seq and paired snRNA-seq/snATAC-seq, we show that fate-biased clones show minimal transcriptional differences at the pluripotent stage by standard clustering yet exhibit distinct chromatin accessibility landscapes at lineage-specific cis-regulatory elements. We find that it is not enrichment of an endoderm chromatin signature but rather pre-existing accessibility at mesoderm-specific regulatory regions that is associated with off-target mesoderm generation. Together, these findings provide insight into how identical differentiation signals yield heterogeneous outcomes from seemingly uniform starting populations, reframing the origin of differentiation variability as governed by heritable cell-intrinsic factors.

Our findings extend the concept of non-genetic clonal memory to germ layer specification in human pluripotent stem cells. While clonal memory has been most extensively described in cancer cells^11,39,40^, where clonal heterogeneity underlies differential drug resistance^12,41^, whether pluripotent hESCs themselves harbour pre-existing clonal fate biases prior to germ layer induction has not previously been addressed. A striking feature of the fate biases we observe is their durability: hESCs were barcoded 14 days prior to differentiation onset, during which cells underwent numerous cell divisions and rounds of passaging where clonally related cells were spatially segregated from their sisters and cousins. This temporal persistence argues against cell-cell communication or local neighbourhood effects as primary drivers, instead implicating stable, cell-intrinsic molecular features. When does this clonal coherence break down during differentiation and for how long is clonal memory maintained as extrinsic signals become progressively more instructive? Future experiments extending lineage tracing through more committed endoderm derivatives, such as pancreatic endoderm, with multiple barcoding timepoints could map bifurcation points where clonal coherence transitions from fate bias to fate commitment.

Chromatin accessibility at cis-regulatory elements can be inherited across cell divisions through histone modification propagation and DNA methylation maintenance^35^, and accessible regulatory regions can be decoupled from immediate transcriptional output, representing poised enhancer states that set up cells for specific responses upon receiving appropriate signals^37,42^. As a number of genes in proximity to DAPs in fate-biased clones are already expressed at the hESC stage, we observed differential regulation of already-expressed genes through differences in chromatin accessibility and motif enrichment. How these accessible chromatin states are maintained, such as through pioneer TF occupancy or propagated through parental histone recycling during DNA replication, cannot be resolved from ATAC-seq data alone. Future single-cell longitudinal chromatin tracking at the clonal level could extend the resolution with which clonal epigenetic dynamics can be interrogated and clarify the nature of regulatory priming in fate-biased clones.

Epigenetic heterogeneity has emerged as an important source of variable differentiation potential in pluripotent cultures^43,44^. While epigenetic memory has been well-characterised in induced pluripotent stem cell (iPSC) reprogramming, where somatic chromatin states persist following cellular reprogramming^45–48^, its role in shaping fate bias within epiblast-derived hESC cultures has been less explored. Our data indicate that a similar principle applies to hESCs, where heritable differences in chromatin accessibility encode fate preferences that manifest following exit from pluripotency. This finding challenges our current understanding of pluripotency and suggests that functional pluripotency at the clonal level may differ from transcriptional pluripotency assessed both globally and at the single-cell level by the expression of canonical markers^49^. WNT3 has been proposed as a predictor of endoderm versus mesoderm differentiation efficiency in hESCs^50^, yet the absence of transcriptional markers at the clonal level in our data highlights the limitations of population-level biomarkers. While an inherent property of pluripotent cells is the capacity to contribute to all cell types of the body, clone-level differences in functional competence may still exist, creating variation both within a cell line and across cell lines. This study presents a snapshot of a single female hESC line in a defined pluripotent stem cell medium undergoing directed differentiation with one protocol. The XCIE subpopulation identified during quality control serves as a reminder that culture adaptation itself represents an additional layer of epigenetic heterogeneity with functional consequences for differentiation. How features of the chromatin landscape compare between male and female cell lines, between hESC and iPSC lines of different tissue origins, and across different culture conditions and extended culture passages remains an important open question with many practical implications for the stem cell field and future clinical applications.

The asymmetric distribution of differential chromatin accessibility observed in fate-biased clones is particularly informative. Mesoderm-biased clones harbour a substantially greater number of DAPs (402 versus 181) and enriched TF binding sites (38 versus 3) than endoderm-biased clones. One model consistent with this asymmetry is that endoderm-biased clones represent a permissive state, where the absence of mesoderm-specific lineage priming allows cells to respond appropriately to TGFβ and WNT signalling, adopting an endoderm fate as intended. In contrast, mesoderm-biased clones possess pre-accessible chromatin at cis-regulatory elements, including regulatory regions near the WNT receptor *LRP6*, where these clones may be hypersensitive to WNT pathway activation by CHIR that diverts them towards a mesoderm off-target despite endoderm-inducing conditions. Although this interpretation is supported by a single experimental system and requires further validation, the broader implication is that if differences in response to signalling are encoded at the clonal level, then fine-tuning differentiation protocols through titration of compounds and growth factors may address only a symptom rather than the root cause of heterogeneity, while strategies such as epigenetic resetting or depletion of off-target-primed subpopulations may instead be required to achieve hESC cultures that are uniformly receptive to targeted differentiation cues.

A central question raised by our findings is whether the fate biases that we observe reflect an *in vitro* culture artifact or a fundamental biological property of pluripotent cells. Epiblast differentiation *in vivo* is spatially and temporally regulated by position-dependent signalling gradients, morphogen fields, and cell-cell interactions that constrain fate choices according to embryonic location^51^. *In vitro* cultures, including those detailed in this study, often lack these spatial constraints, potentially unmasking pre-existing biases that would otherwise be overridden by positional information. In gastruloid formation, individual mouse ESC clones exhibit consistent spatial fate propensities, where developmental precision emerges not from functional equivalence among cells but from the complementary contributions of intrinsically biased clonal populations that collectively ensure robust and reproducible patterning^14^. This raises the intriguing possibility that what appears as an off-target *in vitro* may reflect the operation of biological programmes that serve important functions in the embryo, where different clonal populations are expected to contribute differentially to distinct germ layers. Rather than reflecting a culture artifact, clonal fate biases may reflect a fundamental biological property of pluripotent cells that serves a functional role in the developing embryo, where coordinated action of fate-biased clonal populations may contribute to robust germ layer patterning. This is consistent with the framework proposed by Symmons and Raj^52^, where they describe that what appears as probabilistic variability *in vitro* may represent biologically meaningful diversity-generating mapping that is normally constrained, but not eliminated, by positional information *in vivo*. Our data support a probabilistic model in which fate bias increases the likelihood of a particular outcome without fully committing a cell, and where the duality between memory and plasticity is regulated by the chromatin landscape. The observation that clones transit through transcriptionally indistinguishable primitive streak intermediates yet emerge with distinct fates supports the view that fate divergence is established at the regulatory level prior to transcriptional detection.

The split-well experimental design represents a methodological contribution with broader implications for lineage tracing approaches. By comparing fate outcomes between sister wells derived from the same parental barcoded population, we demonstrate that clonal fate preferences are reproducible across spatially segregated, independently cultured replicates, validating that observed biases reflect intrinsic cell properties rather than microenvironmental variation arising during differentiation. This approach provides statistical power to distinguish heritable fate bias from stochastic variation, and we note that the same biological phenomenon appears weak when analysed within the same sample compared to across replicate samples. Concordance between sister wells was high but not perfect (77% and 69% at hESCs p3 and DE, respectively), and this residual discordance particularly in the hESCs p3 sample where clones would be expected to be most similar captures stochastic sampling effects that disproportionately affect small clones. This underscores the importance of adequate clone sizes for reliable trajectory inference, a consideration for the design of future lineage tracing experiments, particularly those where adjacent or related samples are collected without replicate controls.

Our observations that off-target cell types arising during directed differentiation originate from heritable features across pluripotent culture have implications for stem cell-based cell therapies, meaning that batch-to-batch variation in differentiation efficiency may in part reflect differences in the clonal composition and chromatin landscape of starting cultures rather than variability in protocol execution. While different cell lines may behave differently from each other due to a number of factors, cells within the same cell line may co-exist in pluripotent culture but behave differently in response to signalling cues due to intrinsic heterogeneity. More fundamentally, as high-throughput technologies improve and analysis methods become more sensitive, additional sources of heterogeneity will likely continue to be discovered. The field requires frameworks for distinguishing meaningful heterogeneity from technical noise, and for defining acceptable thresholds of clonal diversity in starting populations for therapeutic applications^1^. Our study presents one example of how off-target populations arise from directed differentiation of pluripotent cultures. Possible strategies to address chromatin-level fate bias include depletion of mesoderm-primed populations through altered pluripotent culture conditions, use of chromatin remodelling agents to reset epigenetic landscapes prior to differentiation, or targeted perturbation of mesoderm-priming transcription factors. Lineage tracing combined with multi-omics provides a powerful approach to systematically dissect the origins of heterogeneity in dynamic biological systems, and the principles identified here are likely applicable beyond endoderm differentiation to other clinically relevant cell types.

## Methods

### Human embryonic stem cell culture

The NOVOe001-A hESC line was maintained in Nutristem (Sartorius: 05-100-1A) medium supplemented with 0.1% penicillin/streptomycin (P/S; Gibco: 15140-122) in tissue culture plates coated with laminin-511 (iMatrix-511; Reprocell: NP892). Medium was changed daily, and cells were passaged every 3-4 days. Passaging was performed by washing cells with PBS (Gibco: 14190-094) and incubated with TrypLE (Gibco: 12563011) for 3-5min at 37°C. Once cells appeared to have reached single cell dissociation after tapping the plate, the reaction was neutralised with Nutristem medium supplemented with 0.1% P/S and 10µM ROCKi (Tocris: Y27632). The cell suspension was centrifuged at 300rcf for 5min, re-suspended in fresh Nutristem medium supplemented with 0.1% P/S and 10µM ROCKi, followed by cell counting using a Nucleocounter (Chemometec: NC-200) and seeded at a density of 15×10^4^ cells/cm^2^. After 24 hrs, the medium was replaced with fresh Nutristem medium supplemented with 0.1% P/S to remove ROCKi. Cells were kept in a humidified incubator at 37°C, 20% O_2_ and 5% CO_2_. Cells were cryopreserved at 1×10^6^ cells per cryovial in StemCellBanker (Amsbio: 11924). hESCs were routinely tested for mycoplasma contamination. The studies outlined in this manuscript were conducted in accordance with ethical guidelines and are covered by the following Danish ethical permits: H-20056277 and H-23039171 (De Videnskabsetiske Komiteer for Region Hovedstaden).

### Lentiviral barcode library generation and delivery

The EF1ɑ-mCherry-ID-16bp-pA library was generated by VectorBuilder using their Barcode Library Construction service (Library ID: Lib241008-1709xgb). For transduction of hESCs with the lentivirus-packaged barcode library, 4×10^3^ cells were seeded on a 48-well CellBIND tissue culture plate (Corning: 3338) coated with iMatrix511 in Nutristem medium supplemented with 0.1% P/S and 10µM ROCKi. The plate was returned to the incubator to allow single cells to adhere to the plate for 1 hr. The lentivirus aliquots were thawed on wet ice and cells transduced by adding the appropriate volume of diluted lentivirus for a multiplicity of infection (MOI) of 4, after which the plate was gently mixed and then returned to the incubator. After 24 hrs, the media was replaced with fresh Nutristem medium supplemented with P/S to remove ROCKi and leftover viral particles. Transduction success was evaluated by flow cytometry based on reporter mCherry expression. The theoretical fraction of uniquely labelled cells was estimated using the approximation, where *N* is the total number of unique barcodes in the library^53^.

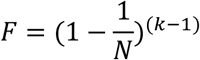

Although MOI 4 was applied, the low number of transduced cells (∼2,400) relative to the library complexity of 40,386,562 unique barcodes (VectorBuilder; Lib241008-1709xgb) yields a theoretical unique labelling rate of >99.99%, confirmed empirically by <1% barcode overlap between independent replicate transductions.

### Definitive endoderm differentiation

hESCs were seeded at 2.4×10^6^ hESCs on 6-well CellBIND tissue culture plates (Corning: 3335) in Nutristem medium supplemented with 0.1% P/S and 10µM ROCKi. 24 hrs after seeding, DE differentiation was induced on cell cultures of ∼80% confluency by replacing medium with RPMI 1640 (Gibco: 61870-044) supplemented with 3µM CHIR99021 (CHIR; Tocris: TB4423-GMP), 100ng/mL Activin A (PeproTech: GMP120-14E) and 0.1% P/S. CHIR was removed following a 24 hr pulse by replacing the medium with fresh RPMI 1640 supplemented with 100 ng/mL Activin A and 0.1% P/S for an additional 48 hrs, until Day 4.

### Flow cytometry

For evaluating transduction success in hESCs, cells dissociated to single cell suspension with TrypLE were incubated with NearIR (Invitrogen: L34976) for live/dead discrimination at 1uL per 1×10^6^ cells in PBS^−/-^ for 15min at RT in the dark, after which cells were fixed with 4% methanol-free PFA (CST: 47746s) for 15 min at RT. Cells were then washed three times in PBS supplemented with 10% KnockOUT Serum Replacement (KOSR; Gibco: 10828-028) and analysed on a CytoFLEX (Beckman Coulter) flow cytometer for expression of mCherry. Data analysis was performed using FlowJo (BD Biosciences).

For isolating barcoded cells based on mCherry expression by fluorescence activated cell sorting (FACS) for subsequent library preparation and scRNA-seq, cryovials containing live cells were thawed at a controlled rate using a ThawStar (Biolife Solutions) and transferred to a wash of RPMI 1640 medium supplemented with 10% KOSR, followed by centrifugation at 300 rcf for 5 min at RT. Cells were then resuspended in PBS supplemented with 10% FBS and 1µL SYTOX 405nm per 1mL of cell suspension for live/dead discrimination, then passed through a 40um Flowmi Cell Strainer (Bel-Art: H13680-0040). Live mCherry positive cells were then sorted using a BD FACSAria Fusion flow cytometer (BD Biosciences) into Protein LoBind tubes (Eppendorf: 0030108116) containing PBS supplemented with 0.04% BSA (Sigma Aldrich: A7284). The cell suspension was then centrifuged at 300 rcf for 5 min at 4°C in a swing-bucket centrifuge and the pellet resuspended in the appropriate volume of cold PBS supplemented with 0.04% BSA to reach targeted number of cells/µL for library preparation input.

### Library preparation for scRNA-seq

Libraries were prepared on cells enriched for reporter mCherry expression by FACS using the Chromium GEM-X Single Cell 3’ Reagents Kit (10X Genomics: 1000691) following the manufacturer’s instructions (CG000731 Rev B), targeting recovery of 20,000 cells per GEM reaction. Libraries were sequenced on a NovaSeq 6000 instrument (Illumina) generating paired-end reads. Successful library preparations were confirmed by Qubit Fluorometric Quantification (Thermo Fisher Scientific) and TapeStation (Agilent). Reads per cell ranged from 31,926 to 42,500 across samples.

### Library preparation for multiome with paired snRNA-seq/snATAC-seq

Single nuclei were isolated from cells enriched for mCherry expression by FACS following the Nuclei Isolation for Single Multiome ATAC + Gene Expression Sequencing protocol (10X Genomics: CG000365 Rev C). Libraries for paired snRNA-seq and snATAC-seq were prepared using the Chromium Next GEM Single Cell Multiome ATAC + Gene Expression Reagents Kits (10X Genomics: 1000283) following the manufacturer’s instruction (CG000338 Rev F), targeting recovery of 7,000 cells per GEM reaction. Libraries were sequenced on a NovaSeq 6000 instrument (Illumina) using separate flow cells for snRNA-seq and snATAC-seq, generating paired-end reads. Successful library preparation was confirmed by Qubit Fluorometric Quantification (Thermo Fisher Scientific) and TapeStation (Agilent). Reads per cell ranged from 42,953 to 98,438 per sample.

### Alignment, data processing and analysis of transcriptomic data

FASTQ files were generated by demultiplexing raw sequencing data using Cell Ranger (10X Genomics). For scRNA-seq samples, reads were aligned to a custom human reference genome (GRCh38, Ensembl release 107) supplemented with the mCherry transgene sequence using Cell Ranger v9.0.0 (cellranger9), generating per-sample cell-by-gene count matrices. For multiome snRNA-seq/snATAC-seq, a custom human reference genome was constructed using cellranger-arc mkref, incorporating the mCherry transgene and its corresponding GTF annotation into the GRCh38 human reference genome. Paired snRNA-seq and snATAC-seq reads were aligned, peak calling performed and processed jointly using Cell Ranger ARC (cellranger-arc), generating per-sample count matrices for both gene expression and chromatin accessibility.

### Barcode retrieval with CellTagR and clonal analysis

Lentiviral barcodes were retrieved from the scRNA-seq and snRNA-seq aligned reads using CellTagR^7,25^ with custom modifications to accommodate our single-barcode library design. BAM files were filtered to retain only reads mapping to the mCherry coding sequence and barcode untranslated region, as well as unmapped reads. Barcode sequences were then extracted from reads containing our construct-specific flanking sequences (5’-ATCGAT-[16 bp]-GAATTC-3’) and their UMI counts were quantified per cell. Extracted barcodes were then collapsed to correct for sequencing errors by clustering similar sequences within a maximum Levenshtein distance using Starcode^54^ default spheres clustering algorithm parameters. Collapsed barcodes were filtered based on a minimum UMI threshold of 2 per barcode per cell, and the resulting count matrix was filtered to exclude cells expressing fewer than 1 or more than 20 unique lentiviral barcodes. Cells were assigned a clonal identity based on pairwise Jaccard similarities of their barcode signatures with a minimum similarity threshold of 0.7. To assess robustness of clone calling to the Jaccard similarity threshold, clones were independently re-called at thresholds 0.6 and 0.8 (compared to the default 0.7) and identical downstream filtering was applied **(Suppl. 2B)**. The validity of clone calling was confirmed by comparing barcode overlap between two independently transduced replicate populations, which revealed <1% barcode collision (14 shared barcodes out of 1,542 unique clones) **(Suppl. 1C, D)**, confirming negligible collision rates within our experimental setup.

### Cell filtering and quality control

Count matrices were imported into R and processed using Seurat v5.4.0^55^ for scRNA-seq and Signac v1.17.1^56^ for paired snRNA-seq/snATAC-seq. For scRNA-seq samples, cells were retained based on the following quality control criteria: a range of 2,500 to 9,500 genes per cell, 3,000 to 65,000 counts per cell, and a maximum of 10% mitochondrial gene counts. For the multiome dataset, nuclei were retained based on 800 to 15,000 genes per nucleus, 1,000 to 150,000 counts per nucleus, a maximum of 10% mitochondrial gene counts, 5,000 to 110,000 ATAC fragments per nucleus, a minimum TSS enrichment score of 1, a nucleosome signal of <1.5, and a blacklist ratio of <0.01. Gene expression data were normalised using scTransform^57^. Cell cycle scores were calculated using CellCycleScoring in Seurat with the canonical S phase and G2M phase gene sets, and cell cycle regression was applied by including S.Score and G2M.Score as variables in scTransform.

### Dimensionality reduction, clustering and cell type annotation

Normalised gene expression matrices for scRNA-seq were used for principal component analysis (PCA) following selection of highly variable genes. Batch correction across samples was performed using reciprocal PCA (RPCA) integration ^58^ as implemented in Seurat using 2,000 highly variable features; this integration was used for visualisation purposes only and all downstream gene expression analyses were performed on the original scTransform-normalised assay. The degree of integration was assessed using the Local Inverse Simpson’s Index (LISI) ^59^. The optimal number of principal components was determined by inspection of an elbow plot, and 50 principal components were selected for UMAP embedding ^60^. Unsupervised clustering was performed by constructing a shared nearest-neighbour (SNN) graph followed by Louvain community detection at resolution 0.2. Cell type annotations were assigned manually to clusters based on expression of canonical lineage markers. For the multiome dataset, chromatin accessibility was processed in parallel using Signac. Raw peak counts were normalised using TF-IDF normalisation and feature selection was performed retaining peaks detected in a minimum of 5 nuclei. Dimensionality reduction was performed using SVD yielding LSI components; LSI component 1 was excluded from downstream analysis following correlation with sequencing depth assessed using DepthCor. UMAP embedding of chromatin accessibility was computed using LSI components 2 to 30. Peak-to-gene linkage analysis was performed using LinkPeaks in Signac, correlating chromatin accessibility at each peak with expression of nearby genes within a 500kb window.

### Clone filtering and fate bias analysis

Following barcode retrieval and clone calling, clones were filtered according to the criteria outlined in **Suppl. 2A**. Clones meeting all criteria and detected at all subsequent sampled timepoints were retained for downstream analysis. Shared clones between same-well and sister-well replicates were identified by exact barcode match and pooled, with non-overlapping clones excluded from replicate comparison analyses.

The clone was used as the primary unit of inference for all clonal analysis, with each clone contributing equally to statistical comparisons regardless of cell number. Clone purity was defined as the fraction of cells within a clone occupying the dominant cluster at a given timepoint. To assess whether observed purity exceeded random expectation, a weighted null model was generated by sampling clone-size-matched cell type assignments from the observed cluster distribution at each stage across 1,000 permutations, and observed purity was compared to the weighted random expectation using a Wilcoxon test. Dominant cluster agreement between same-well and sister-well replicates was assessed by comparing the most abundant cluster identity per clone across replicates and tested against weighted random expectation using a Chi-square test. To assess whether pooling cells across replicate wells altered clone-level purity estimates, purity calculated after pooling same-well and sister-well cells was compared with the mean of purity values calculated independently in each replicate using Pearson correlation. Systematic bias between sampling strategies was evaluated using Bland-Altman analysis. To assess the relationship between clone size and clonal coherence between replicates, Euclidean distance was calculated between the cell type proportion vectors of same-well and sister-well replicates for each shared clone, requiring a minimum of 3 cells per replicate per clone.

A fate bias score was calculated for each clone at the DE stage as the proportion of cells adopting an endoderm fate minus the proportion adopting a mesoderm fate, ranging from −1 (fully mesoderm) to +1 (fully endoderm). Endoderm-biased and mesoderm-biased clones were defined by a minimum of 65% of cells within a clone adopting either a definitive endoderm or early mesoderm identity respectively at the DE stage.

### Differential gene expression analysis

Differential gene expression analysis between cell populations of interest was performed using the FindMarkers function in Seurat with a Wilcoxon rank-sum test, a minimum log2 fold change threshold of 0.25 and a minimum cell detection fraction of 0.1 in either group. Adjusted p-values were calculated using Bonferroni correction. For winner versus loser clone comparisons, analysis was performed on both unmatched and clone size-matched populations to control for clone size effects, where primary clones were downsampled to match the clone size distribution of secondary clones. For endoderm-biased versus mesoderm-biased clone comparisons at hESCs p3 and PS stages, the same parameters were applied.

### Lineage signature scoring

To assess enrichment of lineage-specific transcriptional programmes in fate-biased clones at the hESCs p3 stage, endoderm and mesoderm gene signatures were derived from differential gene expression analysis between endoderm-biased and mesoderm-biased clones at the DE stage. Genes were filtered for *p* < 0.05, where the top 200 genes ranked by log2 fold change were selected for each lineage. Signature scores were calculated using UCell^32^ applied to the scTransform-normalised expression matrix, and differences in scores between fate-biased clone groups at hESCs p3 were assessed using a Wilcoxon test. Gene sets were derived from the same dataset, and results should be interpreted in this context.

### Primitive streak spatial mixing analysis

To quantify the spatial mixing of endoderm-biased and mesoderm-biased clones within the PS subset, a KNN-based mixing index was calculated across varying neighbourhood sizes (k = 10, 25, 50, 75, 100, 150, 200 nearest neighbours) in PCA space using FNN ^61^. For each cell, the fraction of k nearest neighbours belonging to the same fate-bias group was calculated. A mixing index was defined as (observed same-bias fraction − expected) / (1 − expected), where expected was calculated as the sum of squared fate-bias group proportions. Values near zero indicate random mixing and positive values indicate spatial segregation. Statistical significance was assessed by permutation test (100 permutations) in which fate-bias labels were randomly shuffled while preserving group sizes.

### Differential chromatin accessibility analysis

DAPs between endoderm-biased and mesoderm-biased clones at hESCs p3 were identified using the FindMarkers function in Signac with a logistic regression test, a minimum log2 fold change threshold of 0.1 and no minimum cell detection fraction filter. Sequencing depth differences between groups were confirmed to be absent prior to analysis by comparing total chromatin accessibility and TSS enrichment scores between fate-biased clone groups **(Suppl. 4G, H)**. Significant DAPs were defined by *p* < 0.001, log2 fold change > 0.5, and detection in >10% of cells in either group. Nearest gene annotation for significant DAPs was performed using the ClosestFeature function, and a distance threshold of 50kb was applied for downstream target gene analyses. To assess the perdurance of fate-biased chromatin accessibility across developmental stages, DAPs identified at hESCs p3 were compared to DAPs identified at the PS stage using the findOverlaps function in GenomicRanges^62^ with no gap tolerance, and overlap was calculated as the fraction of hESCs p3 DAPs overlapping at least one PS DAP.

### Transcription factor motif enrichment analysis

TF binding motifs were added to the Seurat object using the AddMotifs function in Signac with the JASPAR2020 CORE vertebrate motif database^63^ and the hg38 genome sequence. Motif enrichment in fate-biased DAPs relative to all accessible peaks as background was performed using the FindMotifs function in Signac. Significant motifs were defined by *p* < 0.05 and fold enrichment > 1.5.

### Pathway enrichment analysis

Gene ontology (GO Biological Process, GO Molecular Function) and KEGG pathway enrichment analysis of nearest genes to fate-biased DAPs was performed using enrichR^64–66^ against the GO_Biological_Process_2023, GO_Molecular_Function_2023 and KEGG_2021_Human databases. Terms with *p* < 0.05 were considered significant.

### Statistical analysis and reproducibility

All statistical tests were two-sided unless otherwise stated. Multiple testing correction was performed using Bonferroni correction for differential gene expression analyses and Benjamini-Hochberg correction for motif enrichment analyses. Significance thresholds: *p* < 0.05: *; *p* < 0.01: **; *p* < 0.001: ***; *p* < 0.0001: ****.

## Data and code availability

Raw and processed scRNA-seq and paired snRNA-seq/snATAC-seq data have been deposited to GEO and will be made public at the date of publication. All original code has been deposited to Zenodo and will be made public at the date of publication.

## Acknowledgements

We thank C. Meier and T. Frogne for technical expertise and support with lentiviral barcode library design; and L. Thorén and L.M. Badeby for support with library preparation and sequencing. We also thank C. Honoré, K. Honnens de Lichtenberg, L. Thorén, S. Ulyanchenko, and members of the Morris lab for fruitful discussions and critical comments on this manuscript. This work was financially supported by the Novo Nordisk A/S novoSTAR programme. S.A.M. is supported by grants from the New York Stem Cell Foundation, the Chan Zuckerberg Initiative, and the National Institutes of Health (R35GM153468 and R01DK139580).

## Author contributions

Conceptualization, M.L.-A, N.S.C., S.A.M. and A.L.; methodology, M.L.-A., A.B.G, S.L.H. and Y.S.; investigation, M.L.-A.; formal analysis, M.L.-A and A.B.G.; data curation, M.L.-A and A.B.G.; visualisation, M.L.-A.; writing, M.L.-A, S.A.M. and A.L.; supervision, S.L.H., N.S.C., S.A.M. and A.L.; funding acquisition, N.S.C., S.A.M. and A.L.

