## Supplemental Figures for "Clonal memory in human embryonic stem cells biases fate potential during endoderm differentiation"

**Supplementary Figures**

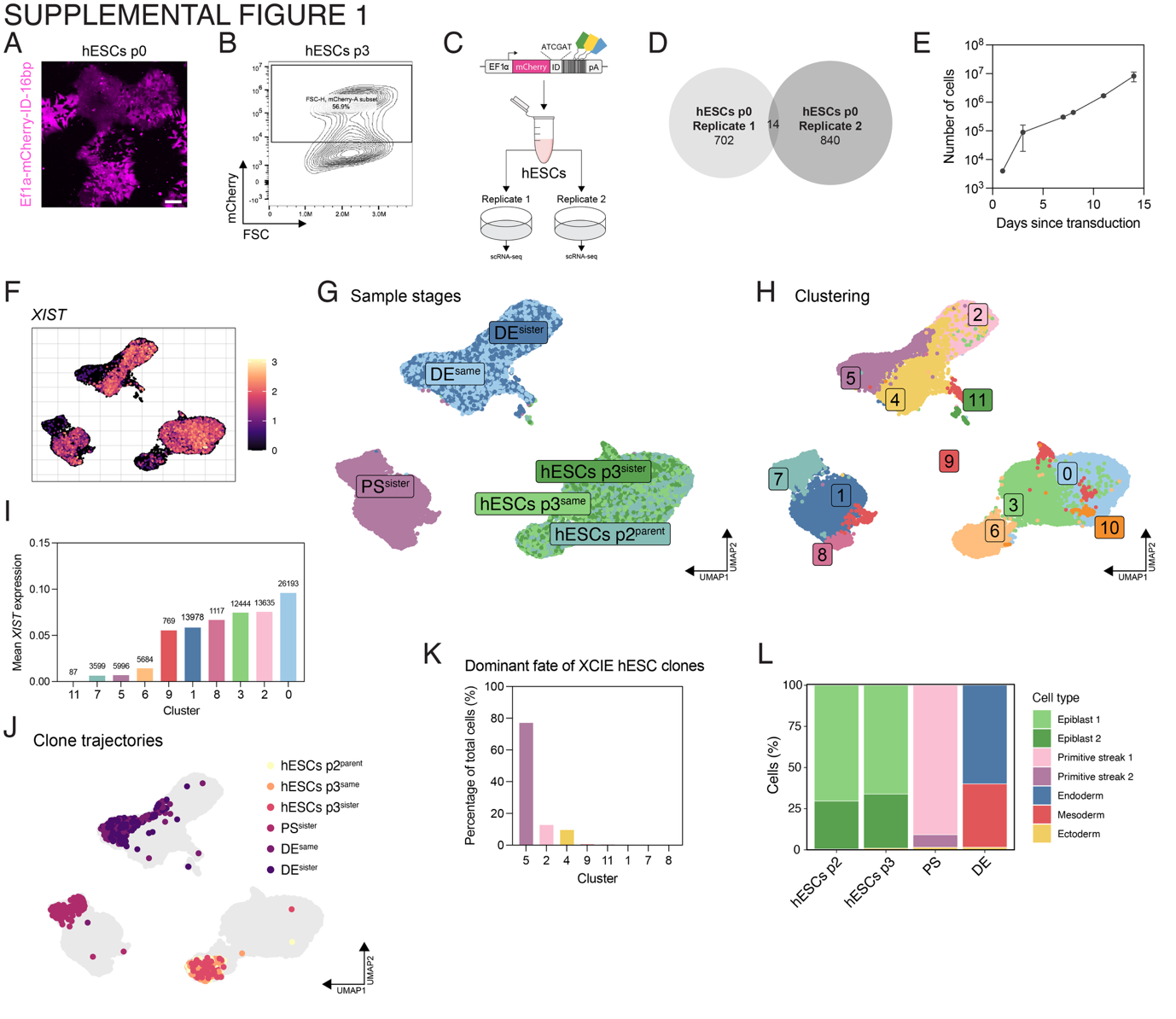

**Supplemental Figure 1:**

**(A)** Fluorescence microscopy of hESCs p0 (7 days post-transduction) showing mCherry expression from the EF1ɑ-mCherry-ID-16bp-pA construct. Scale bar: 100µm. **(B)** Density plot showing flow cytometry gating strategy at hESCs p3 for isolating mCherry expressing cells. **(C)** Schematic of barcode collision validation strategy. Two independent replicate transductions were cultured separately and subjected to scRNA-seq at 7 days post-transduction. **(D)** Venn diagram showing overlap on barcodes between independent replicates (14 shared barcodes out of 702 and 840 unique clones from replicate 1 and 2, respectively; <1% collision rate). **(E)** Cell expansion during pluripotent culture in Nutristem. hESCs (4x10^3^ cells) were seeded after lentiviral transduction (Day 0) and expanded with passaging when confluent. Cells were counted at each passage. Mean ± SD across replicate wells shown. **(F-H)** UMAP dimensional embedding showing **(F)** log-normalised expression of *XIST*, **(G)** timepoint and sampling strategy, and **(H)** unsupervised clustering. **(I)** Mean *XIST* expression across clusters ranked by *XIST* expression level. Sample sizes indicated above bars. **(J)** UMAP dimensional embedding visualising clone trajectory across all sampling timepoints for clones with low *XIST* expression at hESCs p2. **(K)** Fate distribution of XCIE cells at DE (day 4) as a proportion of total cells. **(L)** Stacked barplot showing the proportion of cells assigned to each cluster annotation at each sampled timepoint. Cell type colours as in **Fig. 1D**.

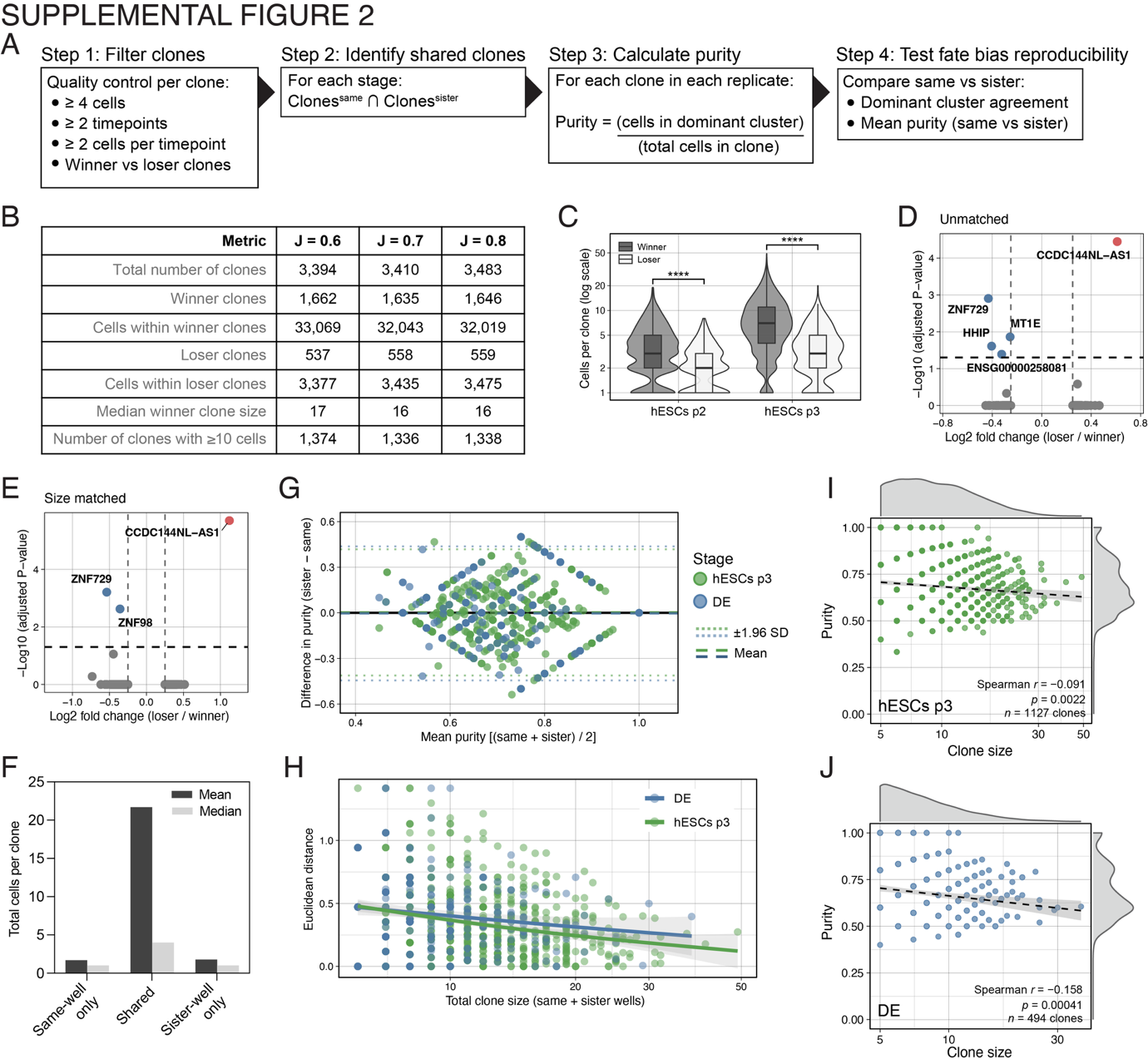

**Supplemental Figure 2:**

**(A)** Pipeline for clone filtering and fate bias analysis. Step 1: Quality control filters requiring ≥4 cells per clone, ≥2 timepoints, and ≥2 cells per timepoint. Step 2: Identification of shared clones between replicates at each stage. Step 3: Purity calculating as a fraction of cells within each clone occupying the dominant cluster/cell type. Step 4: Quantification of fate bias reproducibility by comparing purity and dominant cluster agreement between same-well and sister-well replicates. **(B)** Number of clones and cells per filtering strategy at three different Jaccard similarity thresholds classified as either winner or loser clones. J = 0.7 (default) was used for all main analyses. **(C)** Violin plot comparing clone sizes between winner and loser clones at hESCs p2 and p3. Winner clones are significantly larger than loser clones at both timepoints (*p* < 0.0001, Wilcoxon test). **(D, E)** Volcano plots of differential gene expression between winner and loser clones at hESCs p3. Analysis performed on **(D)** unmatched clones and **(E)** size-matched clones controlling for clone size effects. **(F)** Bar plot of mean and median total number of cells per clone categorised by overlap of replicates with same-well only (clones detected exclusively in same-well replicate), shared (clones detected in both same-well and sister-well replicates), and sister-well only (clones detected exclusively in sister-well replicates). **(G)** Bland-Altman plot related to **Fig. 2K, L** comparing purity measurements between same-well and sister-well. Mean difference ≈ 0 with 95% limits of agreement at ± 1.96 SD. **(H)** Scatter plot showing inverse relationship between total clone size (same-well + sister-wells combined) and Euclidean distance between cells within clones at hESCs p3 and DE. **(I, J)** Clone size versus purity at **(I)** hESCs p3 and **(J)** DE. Purity is the fraction of cells within a clone in the dominant cluster. Marginal densities show distribution of clone size (top) and purity (right). Dashed line is a linear fit with 95% CI.

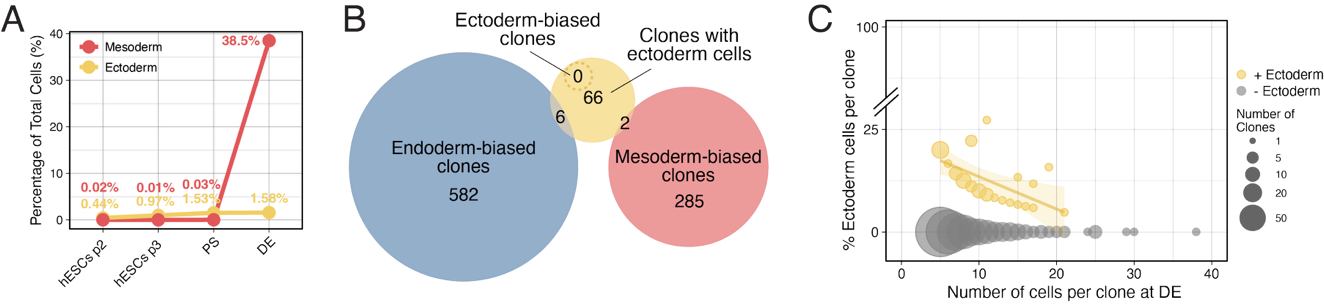

**Supplemental Figure 3:**

**(A)** Line plot showing percentage of total cells classified as the off-target cell type mesoderm (red) or ectoderm (yellow) across developmental stages. **(B)** Venn diagram comparing endoderm-biased and mesoderm-biased clones with clones containing any ectoderm cells at DE. Dashed circle represents ectoderm-biased clones, of which 0 were identified. Minimum number of cells per clone at DE = 5. **(C)** Scatter plot showing the relationship between clone size (number of cells) and percentage of ectoderm cells within each clone at DE. Point size represents the number of clones of a given size; grey points indicate clones without ectoderm cells (92.3%); yellow points indicate clones containing ectoderm cells (7.7%). Yellow line fitted to yellow points.

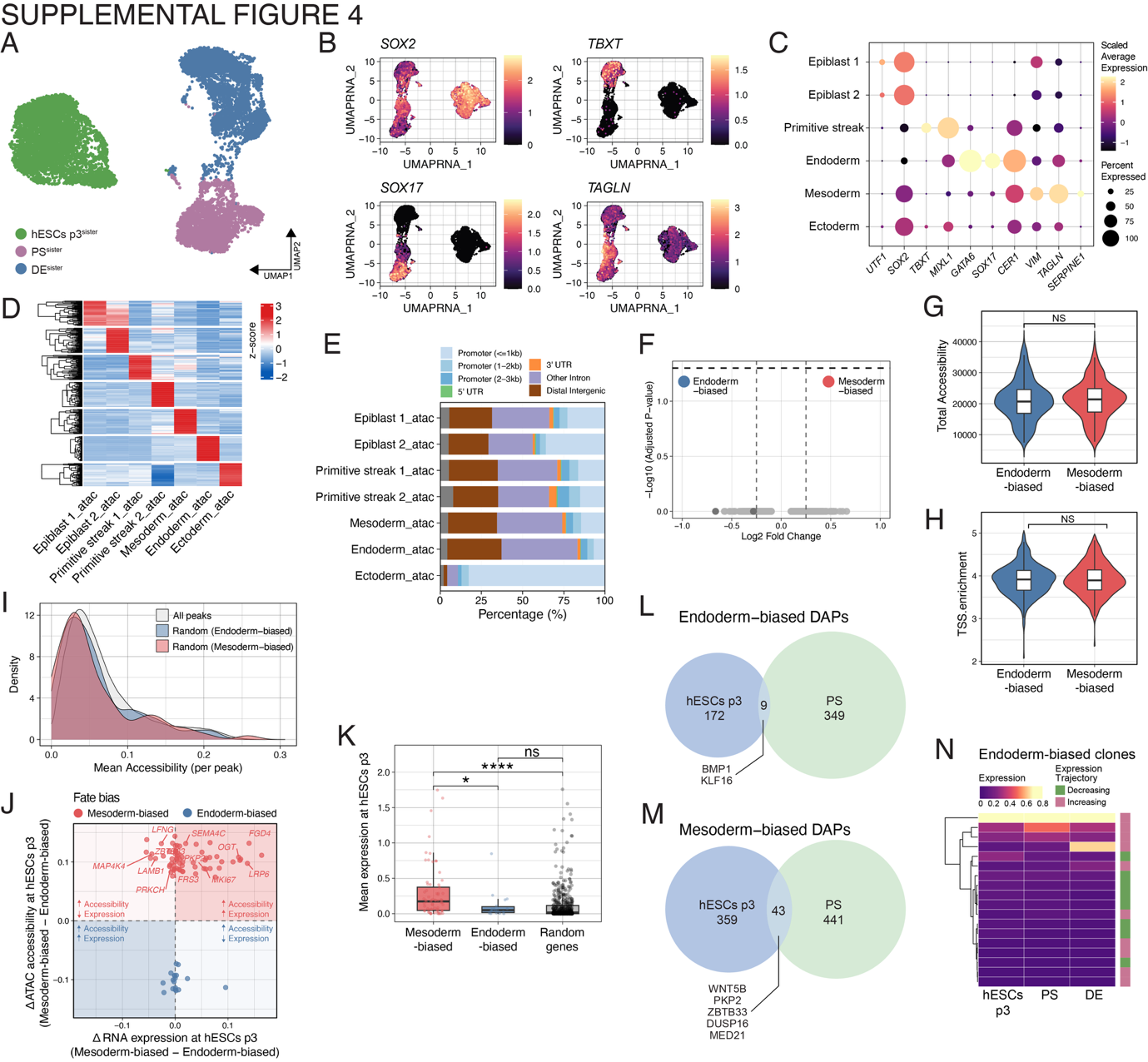

**Supplemental Figure 4:**

**(A)** UMAP dimensional embedding of snRNA-seq coloured by timepoint and sampling strategy. **(B)** UMAP dimensional embedding showing log-normalised expression of indicated markers. **(C)** Dot plot showing expression of lineage-specific marker genes across cell type annotations identified by snRNA-seq clustering. Dot size represents percentage of cells expressing each gene; colour intensity represents average expression level. **(D)** Heatmap of differentially accessible peaks (DAPs) across all cells for each cell type based on ATAC clustering. Rows represent individual peaks hierarchically clustered by accessibility pattern. Colour intensity indicates normalised chromatin accessibility (z-score). **(E)** Stacked bar plot showing genomic distribution of DAPs across snATAC-seq-defined cell types. Peaks are annotated as indicated regulatory elements. **(F)** Volcano plot showing differential gene expression analysis between endoderm-biased and mesoderm-biased clones at hESCs p3. **(G, H)** Violin plots comparing **(G)** total chromatin accessibility (sum of all peak counts per cell) between endoderm-biased and mesoderm-biased cells at hESCs p3 and **(H)** transcription start site (TSS) scores between endoderm-biased and mesoderm-biased cells at hESCs p3. No significant difference was observed in both (NS, Wilcoxon test). **(I)** Density plots showing distribution of mean chromatin accessibility per peak across all peaks in the genome (grey), endoderm-biased DAPs (blue), and mesoderm-biased DAPs (red) at hESCs p3. **(J)** Scatter plot comparing differential chromatin accessibility (y-axis; ΔATAC: mesoderm-biased - endoderm-biased) against differential RNA expression (x-axis; ΔRNA: mesoderm-biased - endoderm-biased) at hESCs p3 for peaks annotated to their nearest gene. Points coloured by fate bias (red = mesoderm-biased DAPs, blue = endoderm-biased DAPs). **(K)** Box plot comparing mean expression of genes nearest to mesoderm-biased DAPs (left), endoderm-biased DAPs (middle), and randomly genomic peaks as control (right) at hESCs p3. **(L, M)** Venn diagrams showing overlap of DAPs between hESCs p3 and PS stages for **(L)** endoderm-biased DAPs and **(M)** mesoderm-biased DAPs. **(N)** Heatmap showing expression trajectories across developmental stages of genes proximal to endoderm-biased DAPs in endoderm-biased clones. Rows represent individual DAP-proximal genes hierarchically clustered by expression pattern. Rows represent individual genes hierarchically clustered by expression pattern. Colour bar indicates expression level (magenta = increasing; green = decreasing).
